# Breaking the Scale: Allometric scaling analysis in Carnivoran families

**DOI:** 10.1101/2021.04.30.442221

**Authors:** Stefanie Navaratnam, Julie Baker Phillips

## Abstract

The analysis of scaling relationships, allometric scaling, has a long history of importance for modelling and predicting biological phenomena. Individual organisms are not truly independent, and as a result phylogenetic corrections are necessary to increase the accuracy of scaling relationships. The relationships between body mass and gestation length have not been previously reported at the Family level, as it was previously thought species had insufficient time to diverge evolutionarily leaving phylogenetic corrections unnecessary. Using a Carnivora supertree, we perform a phylogenetically generalised least squares (PGLS) analysis using life history information largely from the Pantheria dataset. Our results suggest that allometric relationships are maintained in four families: Canidae, Felidae, Herpestidae and Otariidae. Conversely, several evolutionary mechanisms may contribute to the lack of a significant scaling parameter in other families, such as diverse reproductive strategies or positive selection for genes affecting adiposity. In addition, low sample sizes or the inclusion of paternal body masses could alter the presence of significant scaling. Our results suggests that PGLS analyses are informative at the family level, and the absence of scaling can provide insight to understanding of the evolutionary mechanisms that work on the family taxonomic level or below.

**CCS CONCEPTS:** • Applied computing → Molecular evolution; • Computing methodologies → Modeling methodologies.

**ACM Reference Format:** Stefanie Navaratnam and Julie Baker Phillips. 2021. Breaking the Scale: Allometric scaling analysis in Carnivoran families. In *BCB: ACM Conference on Bioinformatics, Computational Biology, and Health Informatics, August 01–04, 2021, Virtual due to COVID-19.* ACM, New York, NY, USA, 7 pages. https://doi.org/10.1145/1122445.1122456

## 1 INTRODUCTION

The analysis of scaling relationships, allometric scaling, is routinely used to model and predict biological phenomena. First performed to investigate the relationship between body mass and metabolic rate [22], it has since been used to uncover the evolution of both morphological and life history characteristics, particularly in relation to body mass [36]. While allometric scaling is often used to understand relationships on the broader scale, it can also reveal intra- or inter-specific evolutionary diversifications in the form of outliers [13].

Historically, ordinary least squares (OLS) analyses were used to demonstrate the statistical relationship between body mass and life history traits such as gestation length, neonate mass and neonate brain weight. OLS analyses assume that data is independent, and does not acknowledge the relatedness of individual organisms. Phylogenetically generalized least squares (PGLS) analyses have been used to not only parameterize the scaling coefficient between two traits, but to elucidate more specific patterns of relationship [8]. For example, the scaling relationship between maternal body mass and gestation length is twice as much when using OLS (M^0.18−0.20^) compared to PGLS regression analysis (M^0.07−0.10^), and subsequent analysis of individual Orders indicated that among closely related taxa, the scaling of gestation is much lower than previously reported with OLS analyses [8]. The contemporary utilisation of phylogenetic-informed statistics necessitates the reanalysis of previously defined relationships and subsequent conclusions.

The scaling relationship of reproductive life history traits is commonly examined, in particular for gestation length in relation to maternal body mass [2, 11, 19]. Past research in Mammalia has operated under the assumption that scaling relationships between traits are generally absent at taxonomic levels lower than the Order due to the close evolutionary relationships. However, many of these originally observed relationships were reported with the use of OLS analyses [28]. PGLS analyses provide a reduced scaling relationship when implemented within Orders, but no direct evidence was provided for its disappearance at taxonomic levels below this, since previous studies had concluded that organisms are too closely related to display a significant scaling relationship [8]. New species-level supertrees with highly resolved phylogenetic relationships can be utilized to correct body mass and gestation data with increasing accuracy. The primary goal of the current work is to examine the scaling relationships at lower taxonomic classifications, particularly within Carnivora, in which myriad studies investigating multiple life history traits exist and are continuously updated.

Carnivora is a diverse Order that includes 286 species divided into 15 families [18, 30]. Prior work from Nyakatura and BinindaEdmonds [30] suggest Carnivora diverged approximately 64.9 million years ago following the Cretaceous-Paleogene boundary. This family encompasses both terrestrial and marine mammals that occur on all continents and vary greatly in terms of the environment they live and their associated life histories. Carnivorans provide an interesting dataset to examine the prevalence of scaling relationships at the family taxonomic level due the the impressive body mass diversity, variety of reproductive strategies, and numerous domestication events, all of which can impact the presence of allometric scaling.

The diversity of organismal mass in Carnivora spans several orders of magnitude, with the smallest carnivores less than 50g (*Mustela nivalis*) and the biggest reaching upwards of 4000kg (*Mirounga leonina*). The diversity of body mass alone among breeds of domesticated dogs is unparalleled by any other land mammal [3].

Two particularly relevant reproductive strategies include delayed implantation (DI), in which prolonged gestation is caused by a cessation of embryonic development [29], and induced ovulation, where eggs are released following behavioral, hormonal or physical stimulation [23]. Embryonic diapause in the form of DI is used by some, though not all, species within a family, or can be completely absent from entire families [10]. Induced ovulation may or may not be present in carnivorans that undergo DI, and this again spans multiple families [17]. Importantly, these adaptations are beneficial in unpredictable settings to maximise fitness and survivability [10]. Reasons for altering reproductive patterns are often environmentally determined, for example due to latitude, predator abundance, population density or home range size [9].

Carnivora encompasses a number of domesticated species. Carnivorans including dogs, cats, ferrets, minks and skunks have all undergone artificial selection for domestication purposes. Numerous studies using OLS have demonstrated that the gestation length of most domesticated species, regardless of the taxonomic classification, does not scale proportionately with body mass intraspecifically [8]. This includes dogs [21], rabbits [40] and horses [4]. Comparatively, cattle breeds displayed significant differences in mean gestation length by species which vary in average dam body mass [5].

In the current work, we seek to elucidate if the scaling relationship between body mass and gestation length is present at the family taxonomic level in Order Carnivora. Here we analyze the relationships using a Carnivoran supertree [30] along with body mass and gestation data [18] to conduct a PGLS analysis, and then discuss possible explanations for the maintenance or absence of scaling relationships within each family, as well as notable outliers within this Order.

## 2 METHODS

### 2.1 Data

Data for adult body mass and gestation length were obtained from the Pantheria dataset [18]. We utilized a Carnivoran supertree for our analyses [30]. Using only species that were present in both the life history data and supertree, we were left with 168 Carnivorans split among fifteen families (Table 1). Four families were too small to be individually analysed, but those species were included for any analysis at the Order level.

**Table 1:** Estimates of Model Parameters for Carnivoran Families.

| Family | n | $\lambda$ | b | $SE_b$ | t | p |
| --- | --- | --- | --- | --- | --- | --- |
| All | 168 | 0.811 | 0.152 | 0.021 | 7.186 | $2.155 \times 10^{-11}$ |
| Canidae | 26 | 0.776 | 0.049 | 0.021 | 2.331 | 0.029 |
| Felidae | 28 | 1.000 | 0.057 | 0.019 | 2.953 | 0.007 |
| Herpestidae | 11 | 0.180 | 0.133 | 0.049 | 2.714 | 0.024 |
| Mustelidae | 34 | 1.000 | 0.206 | 0.058 | 3.573 | 0.001 |
| Otariidae | 12 | 0.694 | 0.152 | 0.072 | 2.110 | 0.061 |
| Phocidae | 18 | 0.000 | -0.018 | 0.033 | -0.546 | 0.593 |
| Ursidae | 8 | 0.000 | -0.273 | 0.323 | -0.844 | 0.431 |
| Viverridae | 7 | 0.000 | 0.134 | 0.095 | 1.403 | 0.219 |
| Eupleridae | 5 | 0.000 | 0.018 | 0.055 | 0.327 | 0.765 |
| Mephitidae | 5 | 0.000 | 0.212 | 0.114 | 1.863 | 0.159 |
| Procyonidae | 7 | 1.000 | 0.070 | 0.126 | 0.555 | 0.604 |

### 2.2 PGLS

We employed a phylogenetic generalised least squares regression (PGLS), which assumes a Brownian motion model of evolution [37], to estimate the scaling coefficient between log-transformed adult body mass and log-transformed gestation length. Data for the gestation lengths is the time between fertilisation and parturition [38], which does not include the duration of DI (see subsequent subsection for additional information).

All statistical tests were performed in R 4.0.4 using the packages ape [33], caper [31], nlme [34] and phytools [35]. Pantheria data was filtered for species present in the Carnivoran supertree with the best estimate of divergence times. Filtered data was combined with the relevant tree and a PGLS analysis was conducted using the caper package [31]. An analysis was performed at the Order level for Carnivora as well as on each of the eleven families in this order (Table 1). Four families had less than five species, Ailuridae, Hyaenidae, Odobenidae and Nandiniidae, and were removed from further analysis.

### 2.3 Pagel’s Lambda

Pagel’s *λ* is a value that quantifies how closely related species are; i.e., the degree that phylogeny plays in determining the relatedness of species [32, 37]. This value can range from 0 to 1; *λ* = 0 suggests no phylogenetic signal between the traits such that closer relatives are no more similar than distant relatives, and *λ* = 1 claims there to be a strong phylogenetic signal where close relatives are more similar than distant relatives [20]. This value can be chosen when performing the PGLS analysis, or it can be estimated based on the maximum-likelihood approach. For our analysis, we estimated *λ* using a maximum-likelihood approach.

### 2.4 Delayed Implantation

Gestation length data provided in Pantheria does not account for the DI period of some species, which influenced the presence of significant scaling relationships at the family level. In families with DI, we performed a PGLS analysis including the DI period [17]. Of major interest was Mustelidae which contains an even split of species with and without DI (n = 13 and n = 11 respectively). A separate PGLS was performed between body mass and gestation length for mustelids who experience DI, as well as for those who do not. *λ* was generated by the maximum-likelihood approach.

### 2.5 Sexual Dimorphism in Body Mass

The Pantheria dataset includes a mix of male and female body masses. Sexual size dimorphisms are widespread throughout Carnivora [23, 39], thus potentially skewing gestation analyses in favour of the male body mass. Two families, Ursidae and Phocidae, were analyzed to investigate the effect of correcting paternal to maternal body mass. Within Ursidae, only the body mass of the outlier *Ursus maritimus* was altered to a female body mass. In Phocidae, all paternal body masses were replaced.

A preferred method would be to use the sexual size dimorphism (SSD) ratio on the male body masses listed in Pantheria to generate the female body masses, though records for Phocidae do not contain enough data to make this a feasible analysis. In addition, SSD ratios vary across papers [9, 12]. Instead, values for female body masses were obtained from various sources (*Monachus schauinslandi*, *Monachus monachus*, *Mirounga angustirostris*, *Mirounga leonina*, *Leptonychotes weddelli*, *Ommatophoca rossi*, *Hydrurga leptonyx*, *Cystophora cristata*, *Erignathus barbatus*, *Halichoerus grypus*, *Phoca groenlandica*, *Phoca fasciata*, *Phoca largha*, *Phoca caspica*, *Phoca sibirica*, *Phoca hispida*, *Phoca vitulina* female masses obtained from [25]; *Lobodon carcinophaga* from [1]). A PGLS analysis was performed on log-transformed female adult body mass and log-transformed gestation length to determine the impact of paternal and maternal body mass on scaling ratios.

## 3 RESULTS

Taxonomic classifications below the Order level are assumed to possess organisms that are too closely related to display a significant scaling relationship [8, 28]. While some of our results contest this assumption (Table 1), there is legitimacy in this statement in that species within some families have not had enough time to diverge evolutionarily to produce a significant correlation. However, every species represents an independent trial that relies not on phylogeny [9]. In accordance with this, *λ* = 0.0000010 for six Carnivoran families in our analysis, indicating a weak phylogenetic signal (Table 1). In this instance, when a phylogenetic signal is barely detectable, results from the PGLS are equivalent to an OLS analysis, as predicted (data not shown; [37]).

### 3.1 Families Maintaining Scaling Relationship

PGLS analysis performed on all species available in Pantheria within Order Carnivora (n = 168) displayed a highly significant relationship (*p* = 2.155 10−11) between log-transformed adult body mass and gestation length, similar to previously published values (Figure 1) [8]. *λ* set to maximum likelihood generated *λ* = 0.811, indicating a strong phylogenetic signal.

**Figure 1:**
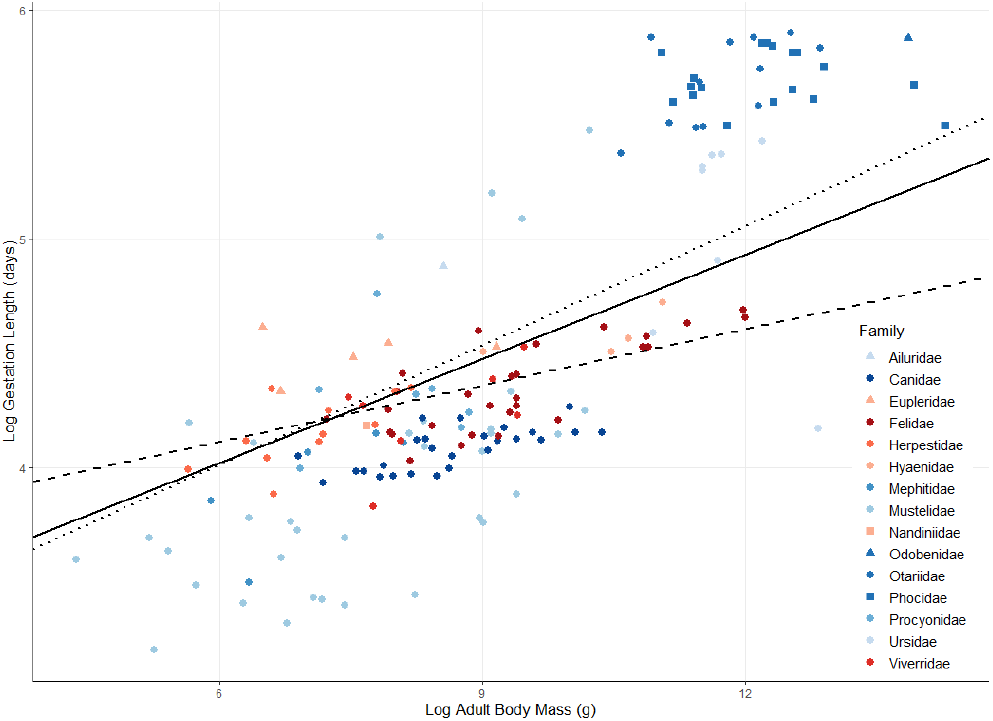
Body mass and gestation scales at the order level using PGLS. Logarithmic plot of gestation length (days) against adult body mass (g) for species in Carnivora. Feliform species are represented by red circles, while caniforms are represented by blue circles. Families of the same colour but different shape are grouped by sister taxa. Solid line represents the phylogenetically corrected linear model for the entire order (*n* = 168, *b* = 0.152, SE = 0.021, λ = 0.811, n = 2.155 10−11) while the dashed line represents that for Feliformia (*n* = 56, *b* = 0.082, *SE* = 0.017, *λ* = 0.991, *p* = 1.698 10−5), and the dotted line for Caniformia (*n* = 112, *b* = 0.174, *SE* = 0.030, *λ* = 0.825, *p* = 6.749 10−8). Mass and gestation length data were taken from the Pantheria database [18] and was linked to a Carnivoran supertree [30].

The PGLS analysis performed on each of the 15 families of Carnivora suggest that three maintain a significant scaling relationship, Felidae (*n* = 28, *n* = 0.007), Herpestidae (*n* = 11, *n* = 0.024), and Canidae (*n* = 26, *n* = 0.029) while Otariidae was very close to significance (*n* = 12, *n* = 0.061) (Table 1, Figure 2). Setting *λ* = 1 in all family analyses assumed a full phylogenetic signal and did not affect the presence of significant scaling relationships (data not shown).

**Figure 2:**
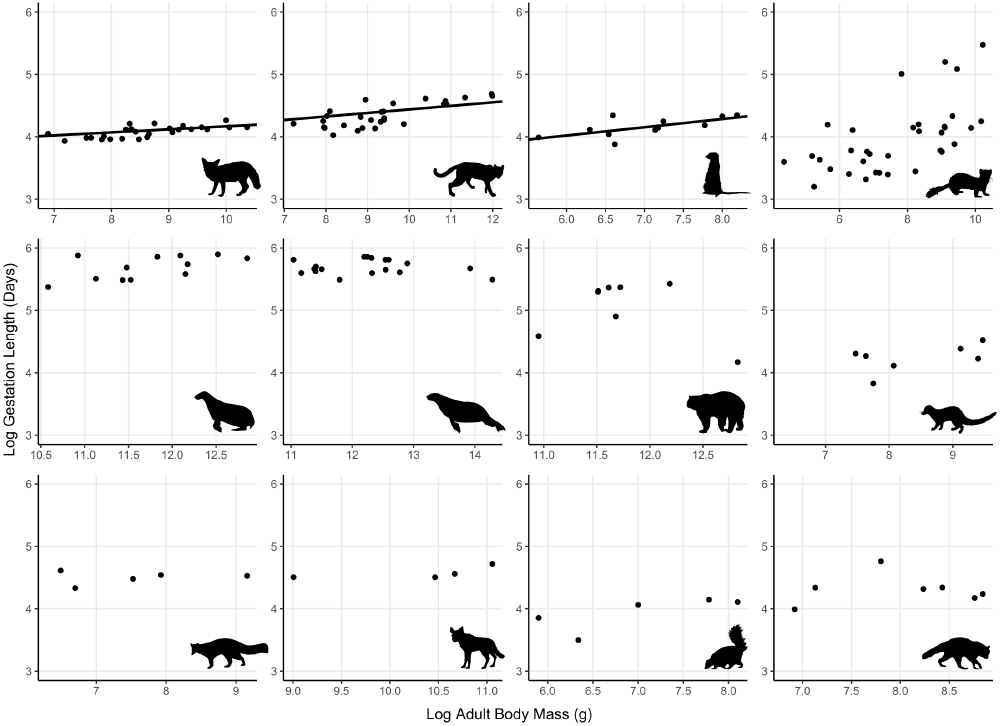
Body mass scales with gestation length in three of the 15 Carnivoran families. Logarithmic plot of gestation length (days) against adult body mass (g) for species within each family of Carnivora. Families shown from left to right, top to bottom are Canidae, Felidae, Herpestidae, Mustelidae, Otariidae, Phocidae, Ursidae, Viverridae, Eupleridae, Hyaenidae, Mephitidae and Procyonidae. Ailuridae, Nandiniidae and Odobenidae are omitted as all have one species each. Mass and gestation length data were taken from the Pantheria database [18] and was linked to a Carnivoran supertree [30]. Images taken from PhyloPic.

### 3.2 Delayed Implantation

Data on the body mass and gestation length of Family Mustelidae was independently analysed to determine if DI had an impact on the scaling relationship. Mustelidae constituted the largest family within the Pantheria dataset (*n* = 34). PGLS analysis between logtransformed adult body mass and absolute gestation length (i.e., not including DI length), demonstrated a highly significant scaling relationship (*P* = 0.001, Figure 2). Species were separated by the absence (n = 11) or presence (n = 13) of DI as reported in the literature [17]. Subsequent PGLS analyses between log-transformed adult body mass and gestation length including DI length revealed that the former group maintained a significant relationship between body mass and gestation length (*p* = 0.0005, *λ* = 0), while the latter did not (*p* = 0.314, *λ* = 0) (Figure 3). Removing two outliers from the group of mustelids with DI (*Mustela lutreola* and *Neovison vison*) remained insignificant (*p* = 0.397) and generated a negative slope (*b* = −0.0431).

**Figure 3:**
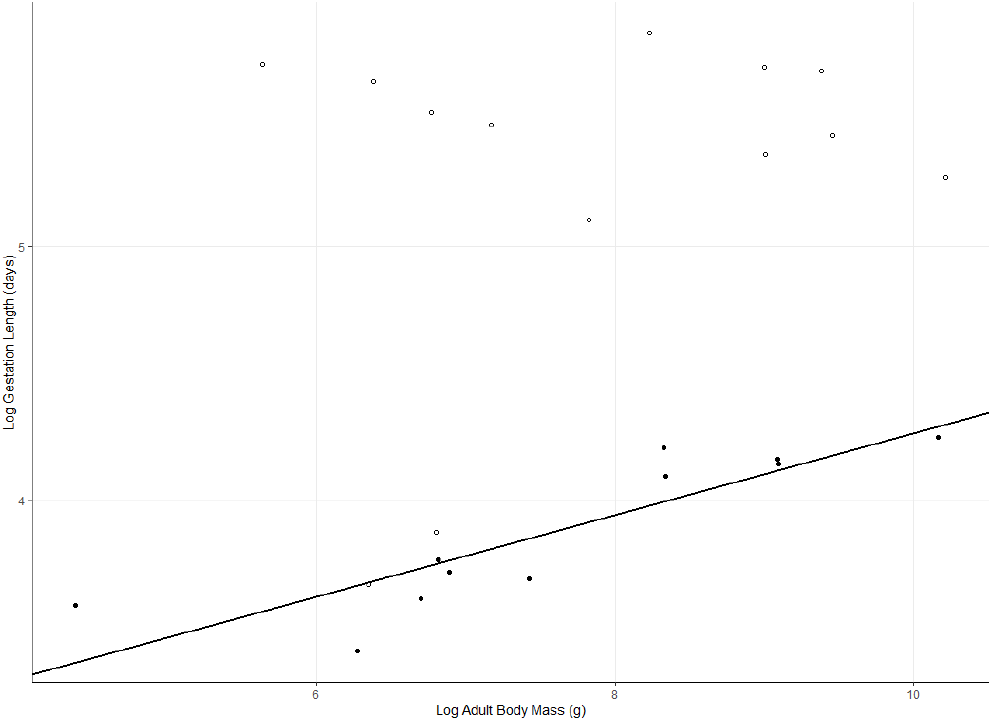
Delayed implantation affects the scaling relationship in Mustelidae. Logarithmic plot of gestation length (days) against adult body mass (g) for species in Mustelidae (n = 24). Circles represent species with DI, while filled circles represent those without. Mustelids without DI exhibit a significant scaling relationship with a scaling coefficient b = 0.161 SE = 0.031). Mass and gestation length data were taken from Heldstab et al (2018) and was linked to a Carnivoran supertree [30].

### 3.3 Sexual Dimorphism in Body Mass

Observations of the Pantheria dataset revealed the inclusion of male body masses for some species listed in Carnivora. Sexual size dimorphisms are widespread throughout Carnivora [23, 39], with ratios of male to female body mass reaching anywhere between 0.17 (*Panthera tigris*) and 6.205 (*Mirounga leonina*) [12, 17], thus potentially skewing gestation analyses in favour of the male body mass.

Ursidae was chosen for this analysis because qualitatively a clear outlier was present in the data which was subsequently identified as the polar bear, *Ursus maritimus* (Figure 2). The polar bear has the largest sexual size dimorphism (SSD) ratio within its family (SSD ratio = 2.44, [9]). To determine if inclusion of the female body mass affected the scaling coefficient, the male body mass for *U. maritimus* was replaced with the adult female body mass to estimate the scaling coefficient. Performing the analysis on Ursidae with the male polar bear mass garnered a steeper slope (*b* = −0.273, *SE* = 0.323, *p* = 0.431, *λ* = 0) than when including the female body mass (*b* = −0.099, *SE* = 0.443, *p* = 0.830, *λ* = 0, Figure 4). Removing the polar bear altogether from Ursidae resulted in a sign change in the value of the slope (*b* = 0.637, *SE* = 0.257, *λ* = 0) and simultaneously produced an almost significant scaling relationship (*p* = 0.056, Figure 4).

**Figure 4:**
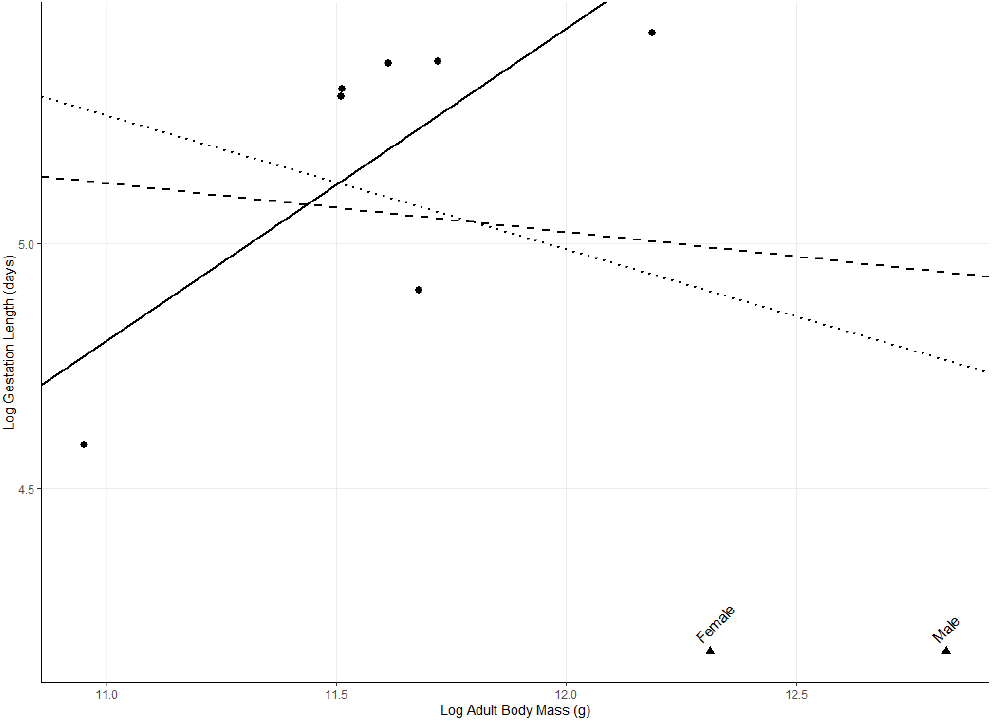
Scaling of Ursidae is constrained by sexual size dimorphism and the presence of an outlier. Logarithmic plot of gestation length (days) against adult body mass (g) for ursids (n = 8). Triangles represent female body mass and male body mass of U. maritimus. Dotted line represents the scaling relationship when using the male mass (b = 0.769, *SE* = 0.444); dashed line represents the scaling relationship when using the female mass (b = 0.253, *SE* = 0.829, *P* = 0.830); solid line represents the scaling relationship when U. maritimus is removed (b = 0.674, SE = 0.198, P = 0.056). Female body mass data obtained from [9].

Phocidae was also analyzed for the impact of sexual dimorphism due to a broad range of SSD ratios (eg, 1.007 for *Leptonychotes weddellii* and 6.205 for *Mirounga leonina*, [12]). In some species, females are larger than males. All of the body masses of Phocidae (*n* = 18) were changed to female body masses if available in the literature (see Methods for all data sources). PGLS analysis revealed that the scaling relationship remained insignificant (*P* = 0.919, *λ* = 0) despite the changes to body mass (Figure 5).

**Figure 5:**
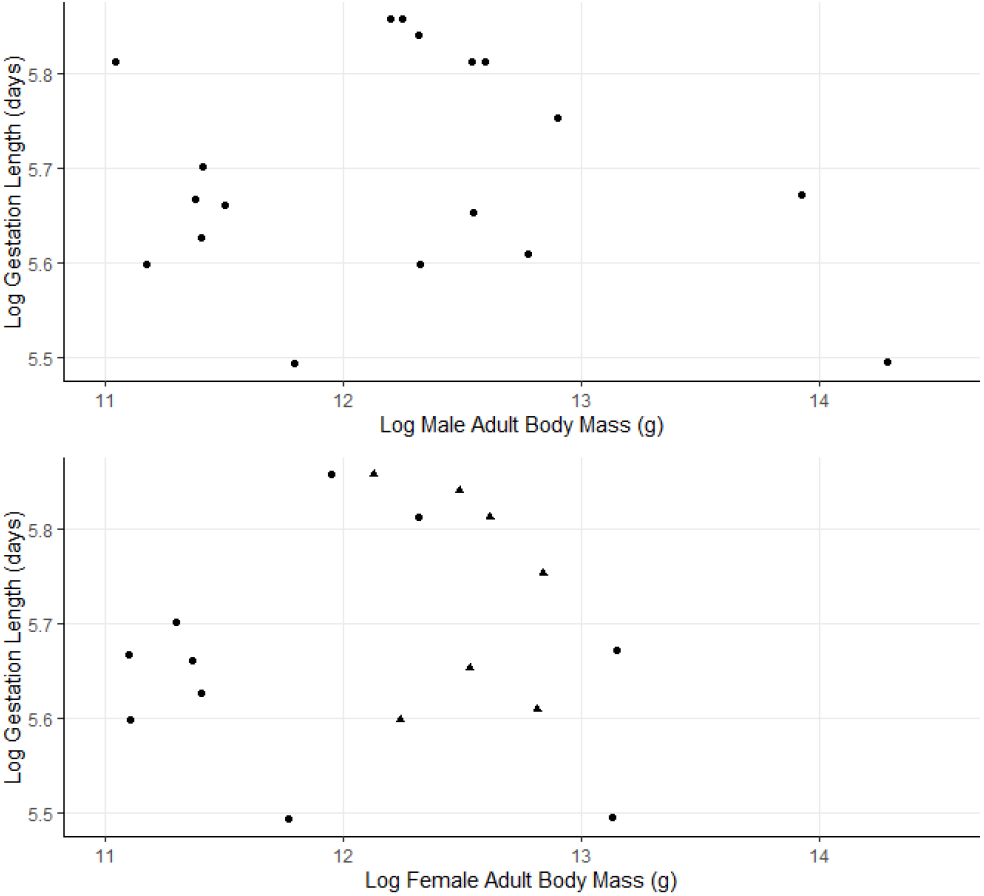
Scaling is insignificant at the family level in Phocidae. (A) Logarithmic plot of gestation length (days) against adult male body mass (g) (*n* = 18). The scaling coefficient is 0.018 (*SE* = 0.033, P = 0.593). (B) Logarithmic plot of gestation length (days) against adult female body mass (g). The scaling coefficient is 0.004 (SE = 0.040, P = 0.919). Triangles represent species in which females are larger than males.

## 4 DISCUSSION

Phylogenetically informed analyses are important and significant at the family level for some families. Our analyses reveal that it is possible for scaling relationships to exist at the family level and that deviations from scaling indicated interesting evolutionary patterns unique to those families within the Order. Data presented here provides evidence that families have had sufficient divergence which result in phylogenetically informed scaling coefficients.

Felidae, Herpestidae, Canidae and Otariidae have few physical commonalities, yet despite the various body plans each family displays a significant scaling parameter. Canids and felids predominantly have intermediate body sizes, otariids are large marine mammals and herpestids are no bigger than 4kg [18]. However from an evolutionary perspective, these four families appear to have diverged most recently out of all the Carnivoran families according to the supertree used in this analysis, with the exception of Hyaenidae whose family was too small to analyze, and Phocidae [16]. In this regard, short divergence times are sufficient to result in significant scaling relationships. Additionally, these four families are of the six largest in Carnivora, perhaps indicating that large families are necessary to demonstrate significance. Alternatively, and maybe equally as important, these families have greater information availability on their species’ gestation lengths (e.g. Viverridae was missing gestation length data for 28 out of the 35 species).

A PGLS analysis adds value over an OLS analysis by revealing the extent to which phylogeny impacts the life history variables estimated. For Felidae, Canidae and Otariidae, the value of Pagel’s *λ* indicates that closely related species are more similar in their gestation lengths than distantly related species. The incredibly low estimate for *λ* in Herpestidae is likely the result of a small sample size, but despite both the small sample size and lack of phylogenetic signal Herpestidae displays a significant scaling relationship. The presence of a strong phylogenetic signal does not necessarily constitute that a scaling relationship will exist, as is the case for Procyonidae and Mustelidae. Conversely, a value of *λ* = 0 is much more likely to be associated with an insignificant scaling relationship. This however is contended by *λ* = 0 for mustelids without DI and Ursidae without *U. maritimus*.

Analyses with insignificant scaling relationships still provide insight, even if the use of PGLS is not necessary. Evolutionary mechanisms have acted on the entire family or within the family in such a way that has inhibited a scaling relationship from existing, which in itself adds value to our growing knowledge of carnivoran evolution.

Alterations in reproductive biology and timing within a family can disrupt the significance of scaling relationships. Mustelids comprise more than half of the total animals that utilise DI to optimise birth timing. This trait is plesiomorphic [38], and our results demonstrate that only mustelids who have evolved without DI display a significant scaling relationship. Consistent with Heldstab et al [17], gestation period was greater in species with DI. This contrasts prior findings that North American terrestrial carnivores with DI had relatively shorter gestation lengths than those without DI [10]. We did not see a variation in the body mass of mustelids with and without DI, which is supported by previous research that determined that body mass of female mustelids is not different in species with and without DI [14], but refuted by Linderfors etal [24]. High seasonality and other environmental factors unique to each species may be a key influence to explain the absence of a scaling relationship amongst mustelids with DI [10, 27].

Outliers within a family can have drastic impacts on scaling relationships. Ursidae is characterised by a low number of species and a non-significant scaling relationship between body mass and gestation length. However, removal of *U. maritimus* provides an almost significant relationship (*p* = 0.056). Population genomics reveal that following divergence of the polar bear from its closest relative the American black bear, *U. americanus*, there was positive selection for genes associated with adipose tissue development, a necessary adaptation for the Arctic Circle [26]. Thus, the body mass augmentation in polar bears without subsequent increases in gestation length is likely to be the causal reason for the lack of scaling in Ursidae.

Marine mammals face unique body constraints in comparison to terrestrial mammals. Pinnipeds do not follow Cope’s rule [7], with evidence pointing to selection for a larger body size. Greater body sizes are favoured as they allow for increased diving depths and oxygen storage capability [7]. Within Pinnipedia, Phocidae does not demonstrate the same significant scaling as Otariidae. Despite relative similarities in gestation length, the smaller body size of otariids limits the amount of energy they can store as fat [6]. Phocids are deep-diving mammals while otariids are shallow divers which reduces oxygen storage requirements, relaxing positive selection for larger body size. Following the evolutionary divergence of Phocidae from Otarioidea in the Oligocene, positive selection for the leptin genes has occured in Phocidae lineage [15, 41]. Leptin satisfies the additional surfactant requirement of phocids who require rapid lung re-inflation following bouts of deep diving, providing a mechanism for increasing oxygen capacity without increasing body size [15]. Serum leptin levels are not an indicator of body fat in pinnipeds [41], but is expressed in the lungs, bone marrow and blubber of phocids [15], indicating that there are perhaps additional physiological functions or tissue specificity of the leptin genes that contributes to its selection in Phocidae only [41].

Overall, the results from our analyses uncover complexities associated with evolutionary allometric scaling relationships. While the presence of scaling is not ubiquitous in Carnivoran families, phylogenetically-informed statistics are warranted at this taxonomic level. Both the presence and absence of significant scaling can reveal informative phylogenetic insights or distinctively frame evolutionary mechanisms that act at this classification or below.

## ACKNOWLEDGMENTS

To J. L. Phillips who provided thoughtful feedback on the manuscript.

## Footnotes

Permission to make digital or hard copies of all or part of this work for personal or classroom use is granted without fee provided that copies are not made or distributed for profit or commercial advantage and that copies bear this notice and the full citation on the first page. Copyrights for components of this work owned by others than ACM must be honored. Abstracting with credit is permitted. To copy otherwise, or republish, to post on servers or to redistribute to lists, requires prior specific permission and/or a fee. Request permissions from. *BCB, August 01–04, 2021, Virtual due to COVID-19* © 2021 Association for Computing Machinery. ACM ISBN 978-1-4503-XXXX-X/18/06. 15.00 https://doi.org/10.1145/1122445.1122456

